# *Pseudomonas aeruginosa* utilises host-derived polyamines to facilitate antimicrobial tolerance

**DOI:** 10.1101/2021.12.15.472801

**Authors:** Chowdhury M. Hasan, Angharad E. Green, Adrienne A. Cox, Jack White, Trevor Jones, Craig Winstanley, Aras Kadioglu, Megan Wright, Daniel R. Neill, Joanne L. Fothergill

**Author notes:** **Correspondence:** Joanne L Fothergill or Daniel R Neill, Department of Clinical Infection, Microbiology and Immunology, Institute of Infection, Veterinary and Ecological Sciences, Ronald Ross Building, 8 West Derby Street, Liverpool, UK, L69 7BE. Equal contributions (Daniel R. Neill, Joanne L. Fothergill).

## Abstract

*Pseudomonas aeruginosa* undergoes diversification during infection of the cystic fibrosis (CF) lung. Understanding these changes requires model systems that capture the complexity of the CF lung environment. We previously identified loss-of-function mutations in the two-component regulatory system sensor kinase gene *pmrB*, in *P. aeruginosa* from CF and from experimental infection of mice. Here, we demonstrate that whilst such mutations lower *in vitro* MICs for multiple antimicrobial classes, this is not reflected in increased antibiotic susceptibility *in vivo*. Loss of PmrB impairs aminoarabinose modification of lipopolysaccharide, increasing the negative charge of the outer membrane and promoting uptake of cationic antimicrobials. However, *in vivo*, this can be offset by increased membrane binding of other positively charged molecules present in lungs. The polyamine spermidine readily coats the surface of PmrB*-*deficient *P. aeruginosa*, reducing susceptibility to antibiotics that rely on charge differences to bind the outer membrane and increasing biofilm formation. Spermidine is elevated in lungs during *P. aeruginosa* infection in mice and during episodes of antimicrobial treatment in people with CF. These findings highlight the need to study antimicrobial resistance under clinically relevant environmental conditions. Microbial mutations carrying fitness costs *in vitro* may be advantageous during infection, where host resources can be utilised.

## Introduction

*Pseudomonas aeruginosa* is a ubiquitous environmental bacterium and a metabolically versatile opportunistic pathogen, responsible for severe, acute nosocomial infections (1, 2). It is also the most frequently recovered pathogen of the cystic fibrosis (CF) lung (3), where it causes chronic infection that is associated with pulmonary exacerbations and declining lung function. Such infections are difficult to treat, due in part to intrinsic antimicrobial resistance of *P. aeruginosa*, together with adaptive resistance mechanisms induced by the presence of antimicrobial agents or other environmental factors (4). The chronic nature of *P. aeruginosa* infection of the CF lung necessitates long-term, high dose antimicrobial therapy, creating conditions conducive to the emergence and selection of acquired resistance mechanisms (5). Phenotypic flexibility and a large genome encoding complex regulatory machinery makes chronically colonised *P. aeruginosa* a challenge to eradicate and necessitates frequent review of treatment regimens for people with CF.

Lipopolysaccharide (LPS) is a key component of the Gram-negative outer membrane and can be stabilized by the addition of divalent cations, including Mg^2+^ and Ca^2+^. Cationic antimicrobials, including polymyxin B, colistin and host-derived peptides such as LL37, exert their effects via disruption of cell membrane integrity, but rely on charge differentials with the outer membrane in order to bind (6). Modifications of LPS that reduce cationic antimicrobial binding affinity and penetration can result in resistance (7). One such modification is the addition of positively charged 4-amino-4-deoxy-L-arabinose (L-Ara4N) to the lipid A component of LPS (8), mediated in *P. aeruginosa* by the proteins encoded by the *arnBCADTEF-ugd* operon (9). Expression of operon genes is regulated by two component signalling systems, such as PhoPQ and PmrAB (10, 11), which are activated under conditions of locally decreased divalent cation concentrations. This ensures that the charge of the membrane can be maintained when Mg^2+^ and Ca^2+^ are limited. PhoPQ- and PmrAB-induced expression of the *arn* operon results in high level resistance to both cationic peptides and aminoglycosides (12). Specific mutations in *pmrAB* have been implicated in polymyxin resistance, via upregulation of both the lipid A deacylase *pagL* and the *arn* operon (13-15).

Expression of genes *PA4775, speE2* (*PA4774*) and *speD2* (*PA4773)* (16), located adjacent to *pmrAB* on the chromosome, is also induced by Mg^2+^ limiting conditions, due to the presence of a PmrA binding site close to *speD2* (6). These genes are involved in the synthesis of the polyamine spermidine. During environmental stress or periods of low cation availability, PmrAB stimulates polyamine synthesis and these coat the bacterial surface, increasing the outer membrane charge and providing protection against both antimicrobial agents and oxidative stress (6).

Polyamines are polycationic hydrocarbons, containing two or more amine groups, and are abundant across all three kingdoms of life. Putrescine, spermidine and cadaverine, the principal polyamines of prokaryotes, have been implicated in growth, iron and free radical scavenging, acid resistance, biofilm formation, protection from the phagolysosome, interaction with components of cell envelopes and antimicrobial resistance (17). However, there is uncertainty regarding the effect, if any, that polyamines have on antimicrobial susceptibility in *P. aeruginosa*. Increased susceptibility to multiple classes of antibiotics was observed when PAO1 was cultured with polyamines, (18) but addition of exogenous polyamines to a PAO1 lacking a functional spermidine synthase (*speE2*) partially protected the outer membrane from polymyxin B (6). Whilst the extent of the role played by polyamines in *P. aeruginosa* growth, virulence and antimicrobial resistance has not been fully determined, it is notable that spermine was found to be elevated in the airways of those with CF and that levels have been reported to decrease during treatment of pulmonary exacerbations (19), whilst those of putrescine have been found to decrease(20).

In a previous study, we identified loss of function mutations in *pmrB* in *P. aeruginosa* isolated from the airways of mice, following experimental infection, and in isolates taken from people with CF (21, 22). These isolates showed enhanced susceptibility to multiple classes of antibiotics. Here, we sought to understand why loss of function *pmrB* mutations might be retained in *P. aeruginosa*, in an environment of prolonged antimicrobial exposure, such as the CF lung. As we had observed altered LPS structure in *pmrB* mutants (22), we hypothesised that host-derived molecules might play a role in stabilising the outer membrane of *P. aeruginosa in vivo*, thereby overcoming the lack of PmrAB-driven modifications of lipid A. Here, we propose that the host cationic polyamine spermidine acts in this way, negating the antimicrobial susceptibility phenotype of *P. aeruginosa* lacking functional PmrB. These findings highlight the need to conduct antimicrobial susceptibility testing under environmentally-relevant conditions.

## Results

### *P. aeruginosa* lacking PmrB show enhanced antimicrobial susceptibility *in vitro* but not *in vivo*

We previously described susceptibility to multiple classes of antibiotics in *P. aeruginosa* with naturally acquired loss of function mutations in *pmrB* and in a *pmrB*-deletion strain of LESB65 (22). To determine whether this susceptibility would result in improved infection outcomes following onset of antimicrobial therapy, we infected mice with LESB65 or a LESB65 mutant lacking *pmrB* (LESB65Δ*pmrB*) and then treated with intra-nasal colistin at 6 and 24 h post-infection. In LESB65-infected mice, colistin treatment led to significant reductions in the number of *P. aeruginosa* recovered from both the upper airways (nasopharynx and sinuses) (Figure 1A) and the lungs (Figure 1B), with four out of eight mice clearing the infection completely. By contrast, colistin treatment did not significantly alter the bacterial burdens recovered from LESB65Δ*pmrB*-infected animals (Figure 1A and B). Consistent with our previous findings, the LESB65Δ*pmrB* strain colonised lungs at a higher density than its wild type parent strain (Figure 1B). We subsequently performed *in vitro* antimicrobial susceptibility testing with bacteria recovered from the infections and confirmed that LESB65Δ*pmrB* retained its susceptibility to colistin *in vitro* (Supplementary Figure 1).

**Figure 1.**
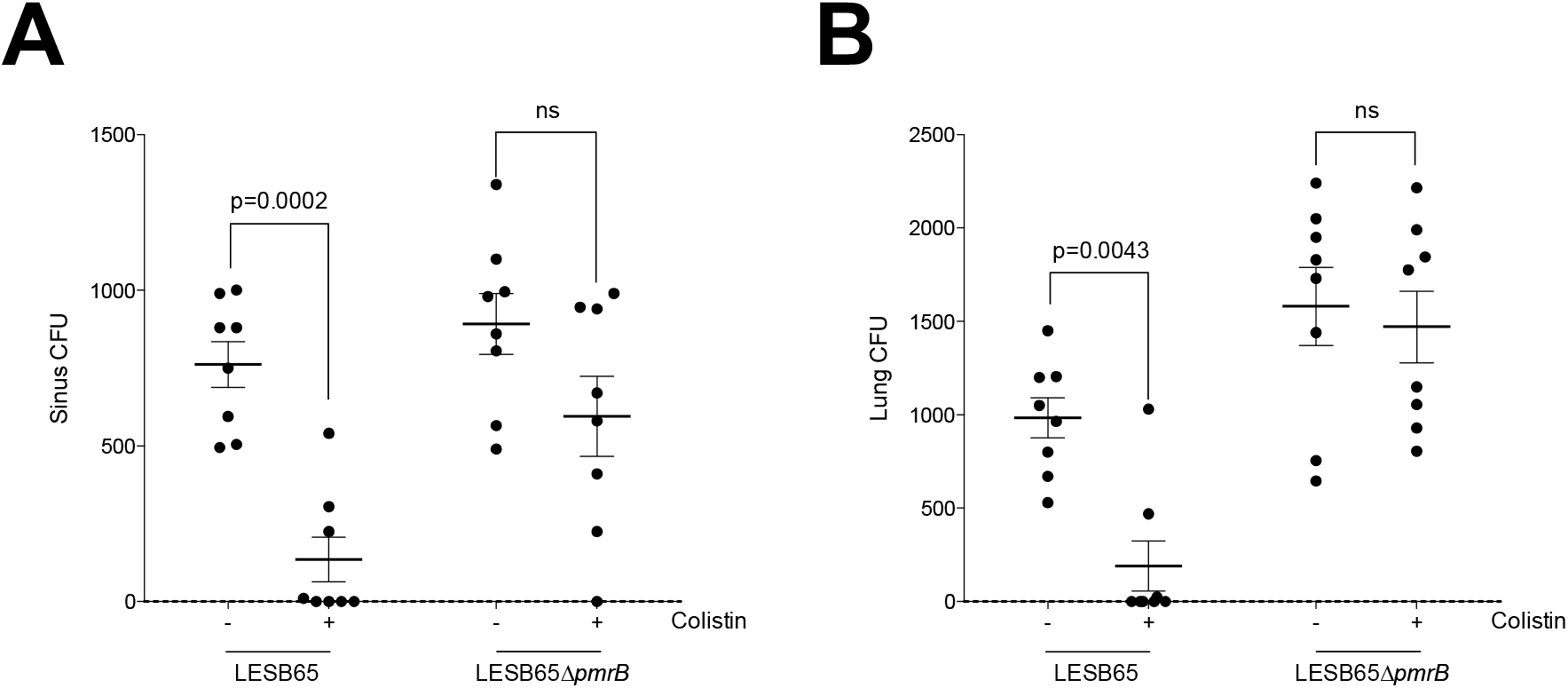
Difference in *in vitro* an *in vivo* antimicrobial susceptibility in PmrB-deficient *P. aeruginosa*. LESB65 or LESB65Δ*pmrB* colony forming units (CFU) in **(A)** sinuses and **(B)** lungs at 48 hours post-infection. Mice were intranasally infected with 2 × 10^6^ CFU *P. aeruginosa*, at 6 and 24 hours post-infection, mice were intranasally administered a 50 μl dose of 20 μg colistin or PBS control. Each circle represents an individual mouse and p values were determined by two-way ANOVA with Bonferroni correction. Data are representative of two independent experiments.

### Proteomics analysis suggests a switch from polyamine synthesis to uptake and utilisation in LESB65Δ*pmrB*

To further explore the environment-dependent antimicrobial susceptibility profile of LESB65 and LESB65Δ*pmrB*, we revisited proteomics data obtained from late-exponential bacterial cultures grown in LB (22). We identified inter-strain abundance differences in a group of functionally related proteins involved in polyamine transport, biosynthesis and metabolism. Bacteria can synthesise polyamines or acquire them via environmental uptake. Genes involved in polyamine synthesis are co-transcribed with *pmrA* and *pmrB* (23) and the proteins encoded by those genes (SpeD2, SpeE2 and PA4775) were found at greatly reduced abundance both in LESB65Δ*pmrB* and in an LESB65-derived isolate with a naturally-acquired missense mutation in *pmrB* (LESB65*pmrB*^L255Q^) (Figure 2A). There is a *pmrA* binding sequence close to the start codon of *speD2* (6) and these data suggest that in the absence of a functional PmrAB system, expression of the operon is significantly reduced. However, the reduced abundance of polyamine synthesis proteins in these strains appears to be offset by a corresponding increased abundance of the polyamine binding, uptake and utilisation proteins of the SpuABCDEFGH operon (Figure 2B). Of the six proteins of this operon that were detected in proteomics analysis, four were significantly more abundant in both LESB65Δ*pmrB* and LESB65*pmrB*^L255Q^, as compared to LESB65.

**Figure 2.**
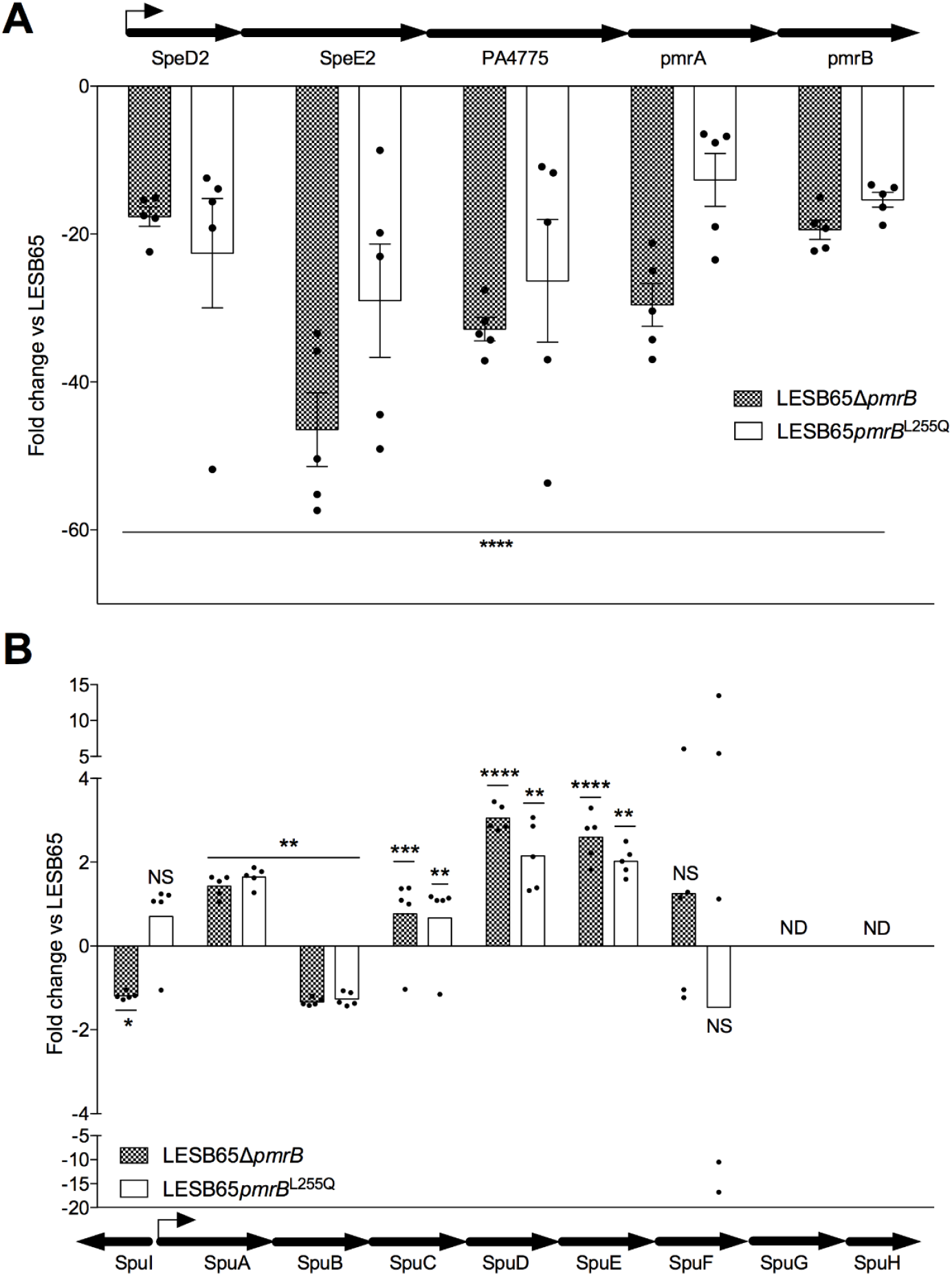
Loss of PmrB influences the relative abundance of polyamine synthesis and utilisation proteins in *P. aeruginosa*. Abundance of proteins of **(A)** the spermidine synthesis operon and **(B)** the polyamine binding, uptake and utilisation operon in LESB65Δ*pmrB* and LESB65*pmrB*^L255Q^ relative to LESB65. Values are fold change vs LESB65 and a negative value denotes decreased abundance. Bars show mean +/- SEM (n = 5 per group) and significance was determined from label free proteomics data using Progenesis QI. NS = adjusted p value > 0.05, * = p < 0.05, ** = p < 0.01, *** = p < 0.001, **** = p < 0.0001, ND = not detected. Block arrows denote gene position and orientation, line arrows show transcriptional start site and directionality. Source data can be found in Bricio-Moreno *et al*. (22), Supplementary Dataset 2.

### Spermidine is abundant in both the airways and increases during infection and antimicrobial treatment

The apparent increase in polyamine binding and acquisition proteins in the *pmrB* mutant strains may be advantageous in environments that are rich in free polyamines. Others have reported polyamine abundance in CF sputum and changes in bioavailability associated with pulmonary exacerbations(19, 20). As the spermidine synthesis proteins were significantly decreased in abundance in PmrB-deficient *P. aeruginosa*, we sought to determine whether the polyamine could instead be scavenged from the environment. We measured spermidine levels in the sinuses and lungs of both uninfected mice and those with chronic *P. aeruginosa* LESB65 infection (Figure 3). Spermidine was detectable at comparable concentrations in sinuses and lungs and was found to increase in the context of infection. This increase is unlikely to result from polyamine production in *P. aeruginosa*, as the levels of spermidine produced by high density cultures of LESB65 or LESB65Δ*pmrB* were ∼1000-fold lower than those detected in respiratory tissues (Supplementary Figure 2).

**Figure 3.**
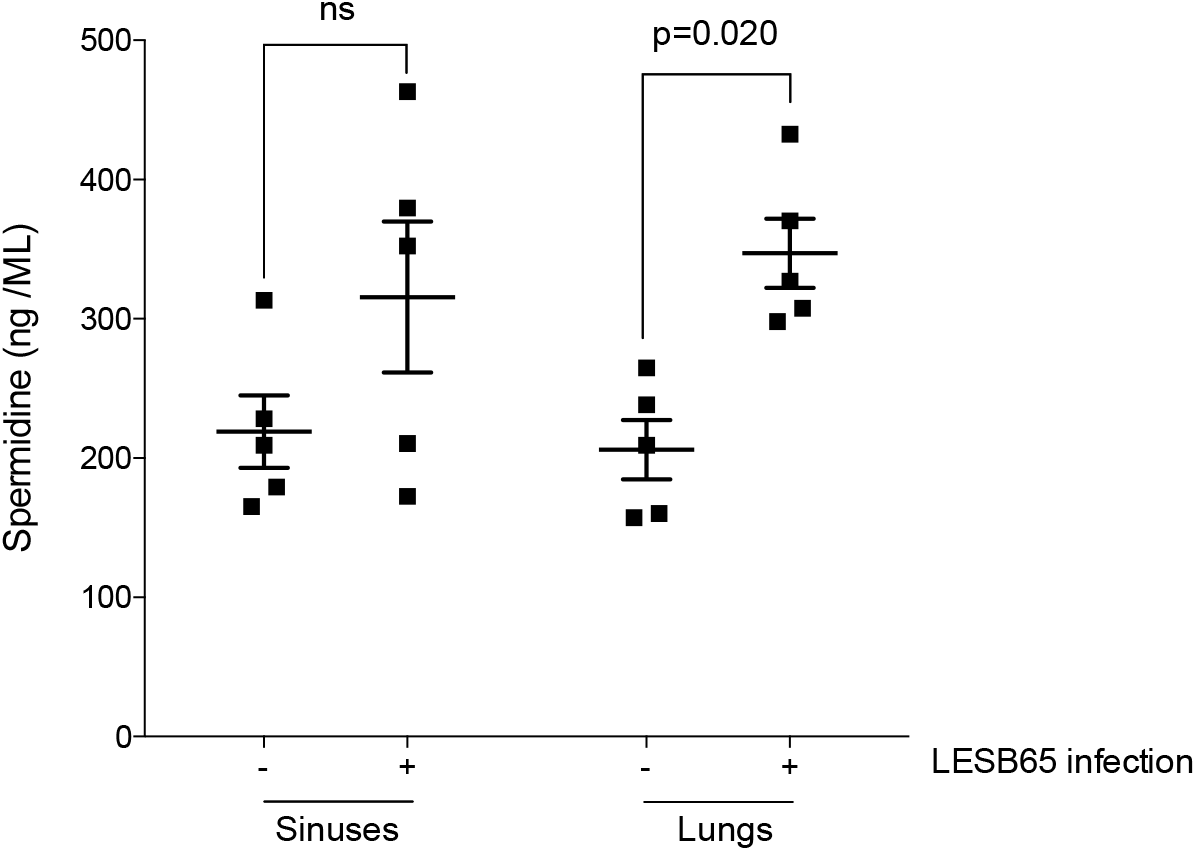
Spermidine is abundant in the murine respiratory tract and bioavailability increases during *P. aeruginosa* infection. Concentration of spermidine in the sinuses and lungs of mice at 48 hours post intranasal administration of PBS (-) or LESB65 (+). Spermidine was measured by ELISA and each square represents a tissue sample from an individual mouse. Significance was determined by two-way ANOVA with Bonferroni correction. Data are from a single experiment.

We also detected free polyamines in CF sputum (Figure 4). Spermidine was quantifiable in samples from 18 of 19 people tested (Figure 4A). There was considerable inter- (Figure 4A) and intra- (Figure 4B) participant variability in sputum spermidine levels, but levels were higher during periods of antimicrobial treatment (Figure 4C), suggesting potential for polyamines to influence treatment efficacy.

**Figure 4.**
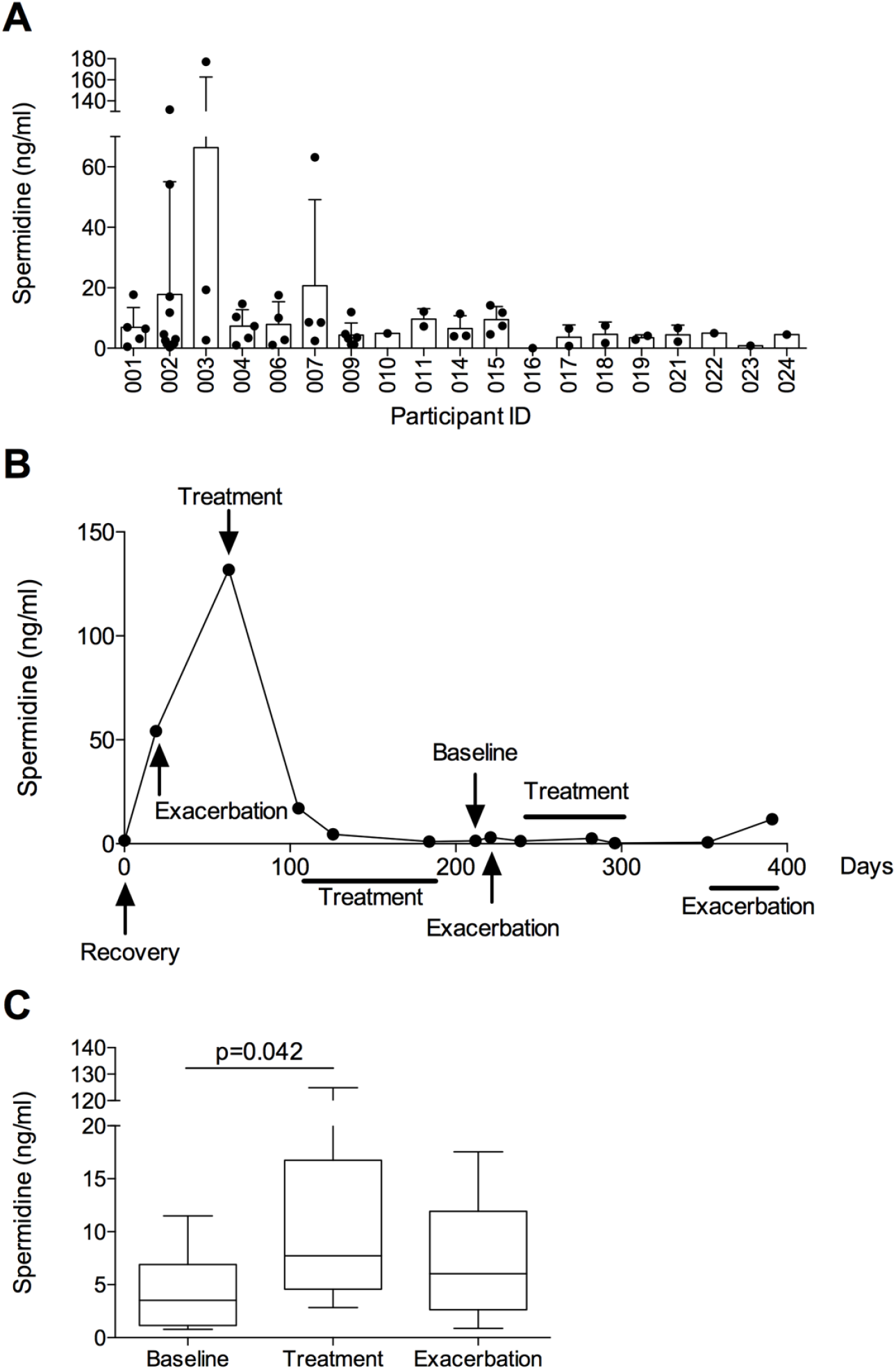
Spermidine is detectable in CF sputum. Concentration of spermidine in CF sputum, determined by competition ELISA. **(A)** Spermidine levels in 19 study participants. Between 1 and 13 samples were available per participant. No two samples from the same participant were collected at the same visit. **(B)** Changes in spermidine abundance in sputum from a single participant over time. **(C)** Collected samples were defined as baseline, treatment or pulmonary exacerbation, defined by participant clinical data. Whiskers show 10-90 percentile. Significance was determined by one-way ANOVA with Dunnett’s multiple comparison testing. Data are from a single experiment.

### PmrB genotype influences *P. aeruginosa* surface interactions with spermidine

Having demonstrated that spermidine is available within the airways, we next sought to characterise whether *P. aeruginosa* might interact with this cationic molecule. Using purified spermidine tagged with fluorescent nitrobenzoxadiazole (NBD), we first investigated whether the polyamine could interact with the bacterial surface and if such interactions were transient or prolonged. For these assays, we used non-toxic concentrations of spermidine, determined by broth microdilution (Supplementary Figure 3). LESB65 and LESB65Δ*pmrB* were co-incubated with 4 mM unlabelled spermidine or spermidine-NBD for 30 minutes before the bacteria were pelleted by centrifugation and resuspended in PBS. We then determined the extent of spermidine binding to the bacterial surface by flow cytometry. Spermidine-NBD bound both LESB65 and LESB65Δ*pmrB*, as evidenced by increasing median fluorescence intensity [MFI] for *P. aeruginosa* co-cultured with labelled vs unlabelled spermidine (MFI 10.4 vs 3.00 for LESB65 with spermidine-NBD vs unlabelled spermidine, MFI 73.3 vs 3.13 for LESB65Δ*pmrB, p<*0.0001 for both strains) (Figure 5). The NBD fluorescence of the LESB65Δ*pmrB* population was significantly higher than that of LESB65 (MFI 73.3 LESB65Δ*pmrB* vs 10.4 LESB65, *p*<0.001). Furthermore, fluorescence was retained longer in LESB65Δ*pmrB* cultures (MFI 73.3 at 0 mins, 67.5 at 30 mins, 62.3 at 60 mins, 56.9 at 120 mins, 54.6 at 240 mins) than in LESB65 cultures (MFI 10.4 at 0 mins, 7.30 at 30 mins, 5.49 at 60 mins, 4.60 at 120 mins, 4.59 at 240 mins) (Figure 5), suggesting prolonged binding or uptake in the LESB65Δ*pmrB* strain. Binding, and surface coating of NBD-spermidine to the *P. aeruginosa* membrane was confirmed by fluorescence microscopy (Supplementary Figure 4).

**Figure 5.**
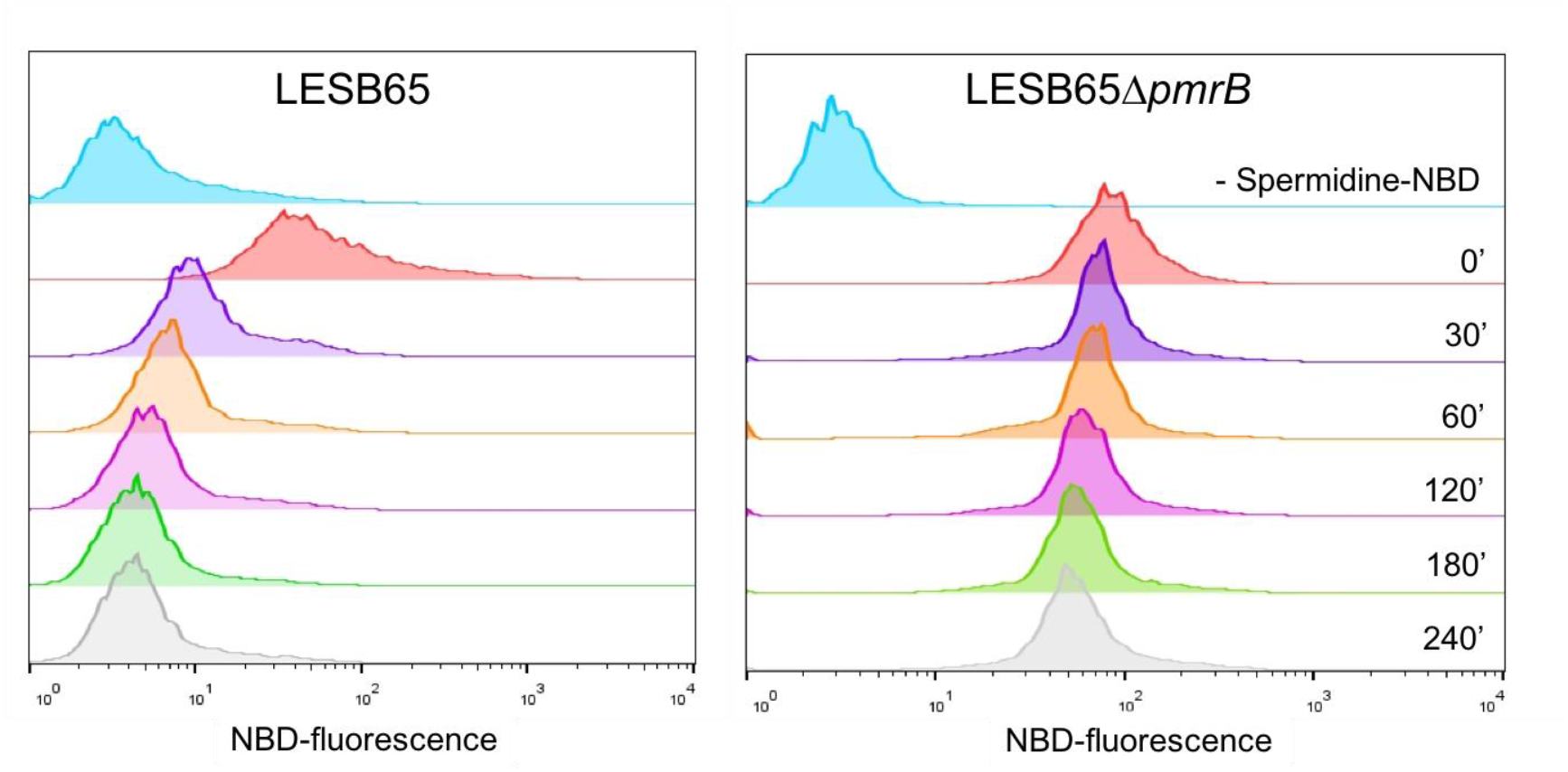
Prolonged interaction with environmental spermidine in PmrB-deficient *P. aeruginosa*. LESB65 and LESB65ΔpmrB from mid-log cultures were incubated for 30 minutes with unlabelled spermidine (blue histogram) or spermidine-NBD (all other histograms), then pelleted, washed in saline and resuspended in polyamine-free PBS. Spermidine-NBD binding to *P. aeruginosa* was determined by flow cytometry at 0, 30, 60, 120, 180 and 240 minutes after co-incubation. Data are representative of two independent experiments.

### Surface-coated spermidine protects *P. aeruginosa* from antimicrobials and offsets the susceptibility associated with loss of PmrB function

Polyamines have been implicated in resistance to several classes of antibiotics, with binding of these positively charged molecules to the *P. aeruginosa* membrane reducing charge interactions with cationic antimicrobials (6, 23). The decreased abundance of SpeD2, SpeE2 and PA4775 in loss-of-function *pmrB* mutants may, therefore, contribute to the observed increases in antimicrobial susceptibility, under conditions where polyamines cannot be readily scavenged from the environment. However, where polyamines are abundant, the increased polyamine-binding potential of LESB65Δ*pmrB* might offset the inherent susceptibility to antimicrobials associated with loss of PmrB function. To explore this idea, we performed MIC assays with *P. aeruginosa* without spermidine, in the presence of spermidine, or with *P. aeruginosa* that had been pre-incubated with spermidine and then pelleted and washed before addition of antibiotics. (Table 1). Assays were performed with PmrB-deficient strains on both the LESB65 and PAO1 backgrounds. In both cases, spermidine increased the colistin MIC50 of the *pmrB*-deficient strain, but not the wild type ancestor, by 4-8 fold. This was the case both when the spermidine was present throughout the assay and when strains were pre-incubated with the polyamine. In polyamine-rich environments, such as the respiratory tract, surface-coating of *P. aeruginosa* with cationic polyamines may achieve a comparable outcome to PmrAB-driven L-Ara-4N addition to LPS lipid A, by increasing the positive charge of the outer membrane. This finding may explain why loss-of-function *pmrB* mutations are retained in *P. aeruginosa* causing chronic infection of the CF lung, despite prolonged, high dose antimicrobial treatment.

**Table 1.** Presence of spermidine reduces antimicrobial susceptibility of PmrB-deficient but not wild type *P. aeruginosa*. The concentration of colistin required to inhibit 50 percent of growth (minimum inhibitory concentration [MIC]50) was determined for LESB65, PAO1 and their PmrB-deficient derivatives. Assays were conducted without spermidine, in the presence of 4 mM spermidine, or with bacteria that had been pre-incubated for 30 minutes with 4 mM spermidine and then washed in PBS before addition of antibiotics. Data shown are the median and range of MIC50 values from 5 independent experiments.

| Strain | Spermidine throughout | Preincubation with spermidine | MIC50 μg/ml [Median (range)] |
| --- | --- | --- | --- |
| LESB65 | - | - | **2** (2-4) |
|  | + | - | **2** (2-4) |
|  | - | + | **2** (1-4) |
| LESB65Δ*pmrB* | - | - | **0.25** (0.125-0.5) |
|  | + | - | **2** (0.5-4) |
|  | - | + | **1** (1) |
| PAO1 | - | - | **0.5** (0.5) |
|  | + | - | **1** (0.5-1) |
|  | - | + | **0.5** (0.25-0.5) |
| PAO1Δ*pmrB* | - | - | **0.125** (0.125-0.25) |
|  | + | - | **2** (0.5-2) |
|  | - | + | **1** (0.5-2) |

### Exogenous spermidine influences *P. aeruginosa* biofilm formation

Polyamines contribute to biofilm formation in both Gram positive and Gram negative bacterial species (24). They serve as structural components of the extracellular matrix, but also stimulate signalling that can promote or inhibit biofilm formation (25, 26). However, the role of polyamines in *P. aeruginosa* biofilm formation is not well described. To determine whether spermidine-induced reductions in antimicrobial susceptibility in LESB65Δ*pmrB* might by augmented by enhanced biofilm formation, we quantified surface attached biofilm by crystal violet staining. The addition of spermidine to cultures had no effect on LESB65 biofilm but led to a ∼2.5-fold increase in LESB65Δ*pmrB* biofilm biomass over a 48 hour period (Figure 6).

**Figure 6.**
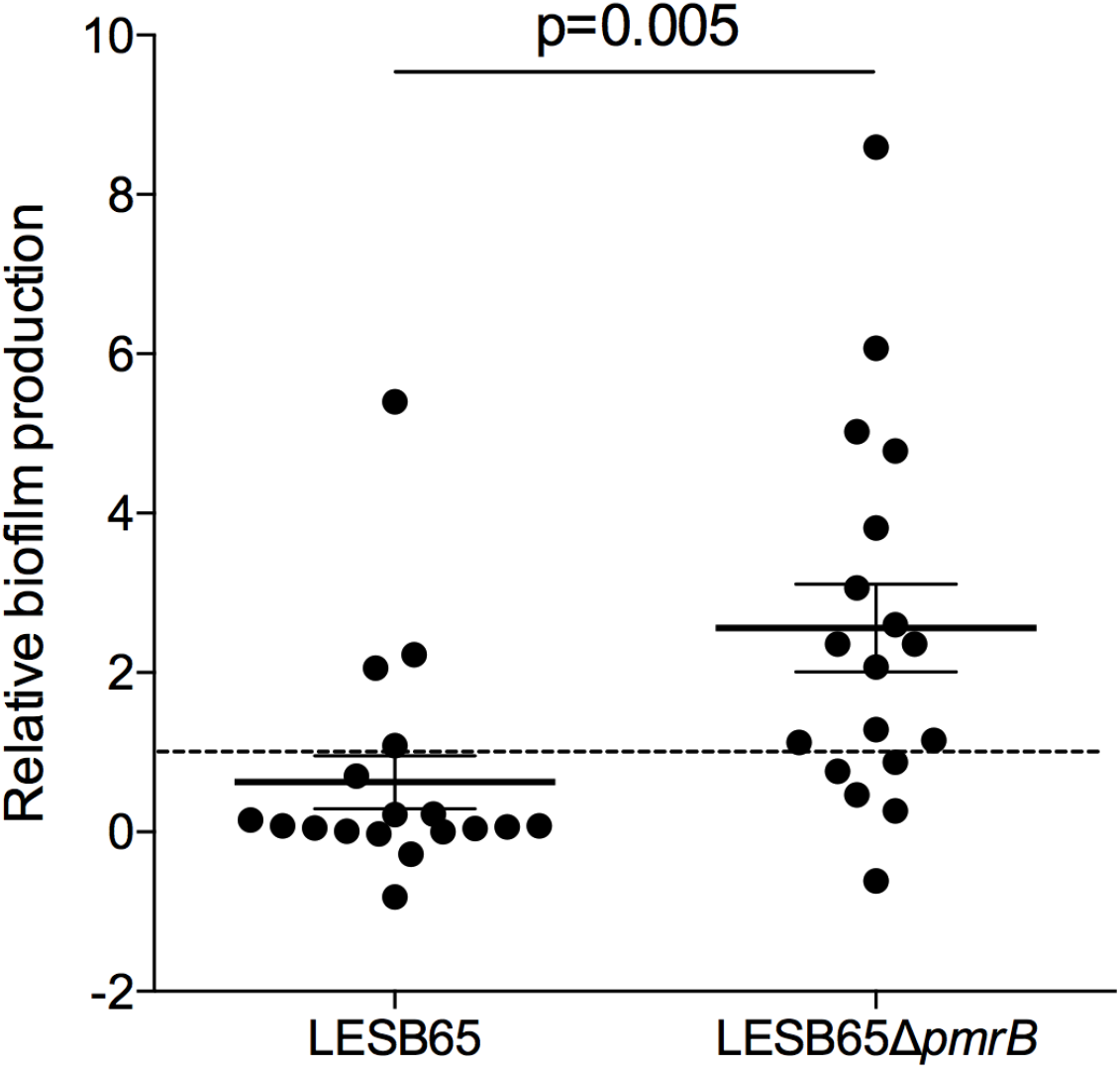
Spermidine promotes biofilm production in PmrB-deficient *P. aeruginosa*. Surface-attached biofilm production was quantified by crystal violet staining, following 48 hours of culture. Data are expressed as fold-changes in biofilm production vs the no spermidine control for each strain. Data are pooled from three independent experiments.

## Discussion

Despite their ubiquity and essentiality across all kingdoms of life, understanding of polyamine biology is limited. *P. aeruginosa* can synthesise polyamines from methionine, arginine or ornithine, but also scavenge them from their environment. The expression of *P. aeruginosa* polyamine synthesis, uptake and utilisation genes is under environmental control, including by the PmrAB two-component regulatory system.

The PmrAB system plays a key role in environmental sensing, stimulating production of polyamines and modification of LPS to buffer the outer membrane charge in conditions of low divalent cation availability (14). Mutations in *pmrB* are frequently identified in *P. aeruginosa* infecting the CF lung (27), and both activating and loss-of-function mutations have been described (13, 22). This apparent dichotomy may reflect environmental differences between individuals or between niches within the host. Where divalent cations or other positively charged molecules, including polyamines, can be co-opted to buffer membrane charge, loss of function *pmrB* mutations may be retained due to the advantages they confer in terms of lysozyme resistance and enhanced adherence to host surfaces (28). However, under conditions where cationic molecules are scarce, activating mutations in *pmrB* would likely confer a greater advantage, by promoting the LPS modifications that reduce binding of antimicrobials to the bacterial outer membrane.

The differences in the kinetics of interaction with spermidine in wild type and PmrB-deficient LESB65 suggest that surface charge may influence the capacity of *P. aeruginosa* to utilise exogenous polyamines. Whilst it is clear that detection of divalent cations through PhoPQ and PmrAB can modulate polyamine synthesis and uptake pathways, as well as inducing surface charge modifications, it remains to be determined whether direct sensing of polyamine abundance can modulate those same processes. An ability to alter surfacecharge, and thus change the efficiency of polyamine binding, in response to the local availability of those molecules, would offer advantages in metabolic resource management.

Similarly, whilst the findings presented here demonstrate spermidine binding to the surface of *P. aeruginosa*, it is unclear whether the changes in antimicrobial susceptibility and biofilm formation that we observed are a direct result of that physical interaction or whether they are a consequence of polyamine-induced signalling. The increase in positive charge associated with spermidine coating of the outer membrane may be sufficient to explain the increased resistance to cationic antimicrobials such as colistin, and spermidine might act as a substrate for biofilm formation, encourage greater surface interactions via the change in membrane charge or aid in chelation of negatively charged biofilm DNA. However, we can’t rule out further contributions from spermidine-induced signalling and others have reported an effect of exogenous polyamines on bacterial pathogen gene expression, including in *P. aeruginosa* (29, 30).

The composition and charge of the outer membrane of Gram-negative bacteria are key determinants of antimicrobial resistance and major barriers to antibiotic uptake (31). The findings presented here go some way towards explaining the dichotomy of retention of loss-of-function *pmrB* mutations in *P. aeruginosa*, in the face of antimicrobial pressure. The susceptibility of PmrB-deficient LESB65 to antimicrobials *in vitro* did not translate to susceptibility within the lung environment and this may be explained by buffering of the negatively-charged outer membrane with host-derived cationic molecules, including spermidine. This highlights the need to perform antimicrobial susceptibility testing under conditions that are relevant to infection and that capture key environmental cues that are sensed by pathogens or with which they interact. This is a particular challenge for those interested in pathogens of the CF lung, given the complexity of that environment and the difficulty in replicating its physical, chemical and microbiological features in the laboratory. However, substantial progress has been made in this area (32), through use of metabolic profiling of CF pathogens (33) and analytical methods that utilise next-generation sequencing data to aid in the benchmarking of new models that aim to replicate conditions of the CF lung (34). As models continue to be refined, consideration should be given to the inclusion of host-derived factors, including polyamines, that might influence pathogen membrane charge and antimicrobial susceptibility.

## Materials and methods

### Bacteria and culture conditions

*P. aeruginosa* Liverpool Epidemic Strain (LES)B65, LESB65Δ*pmrB*, PAO1 and PAO1Δ*pmrB* were used throughout. Deletion of *pmrB* in LESB65 and PAO1 was performed as part of a previous study (22). Bacterial stocks were stored at −80°C in 15% (v/v) glycerol. Prior to experiments, isolates were streaked onto Mueller Hinton (MH) agar and then liquid cultures were prepared in MH broth, unless otherwise stated, from a single colony, and incubated at 37°C in a shaking incubator (180 rpm).

### Sputum samples

Sputum samples were collected from people with CF at the Adult CF centre (Liverpool Heart and Chest Hospital) during periods of stable infection and periods of exacerbation in accordance with ethical approval (IRAS:216408, ethics reference no: 17/NW/0091). Samples were expectorated between 2017 and 2020, and stored at −80°C within 2 h of production.

### Chemicals and reagents

Antibiotics and spermidine were purchased from Sigma-Aldrich (Sigma, UK). Stock solutions were prepared using DEPC water and filtered through 0.22 µm syringe filter.

### LESB65 infection of mice

All infections were performed at the University of Liverpool with prior approval by the UK Home Office and the University of Liverpool Ethics Committee. Female BALB/c mice of 6-8 weeks of age (Charles River, UK) were used for infection experiments and housed in individually ventilated cages. Mice were acclimatised for one week prior to infection. Mice were randomly assigned to an experimental group on arrival at the unit by staff with no role in study design. For infection, 2 × 10^6^ colony forming units of mid-exponential growth *P. aeruginosa* were instilled into the nares of mice that had been lightly anaesthetised with a mixture of isoflurane and oxygen. At 6 and 24 hours post-infection, mice were intranasally administered a 50 μl dose of 400 μg/ml colistin in PBS or else PBS only for control animals, under light anaesthesia. Following this, cage labels were reversed to blind researchers to experimental groups. Mice were culled at 48 hours post-infection and upper airway (sinus and nasopharynx) tissue and lungs were removed post-mortem and homogenised in 3 ml PBS using an IKA T10 handheld tissue homogeniser (IKA, USA). Homogenates were serially dilution onto *Pseudomonas* selective agar (Oxoid, UK) for enumeration of infectious burden. Following enumeration, researchers were unblinded. No animals were excluded from analysis.

### Spermidine ELISA

Spermidine was quantified from lysates of overnight *P. aeruginosa* cultures, from mouse upper airway (sinus and nasopharynx) and lower airway (lung) tissue homogenates, and from CF sputum. Bacterial lysates were prepared by sonication. Overnight cultures were pelleted by centrifugation and resuspended in 1 ml PBS prior to sonication. Mouse tissues were dissected at 48 hours post intranasal administration of 2 × 10^6^ colony forming units of mid-log phase LESB65 (infected group) or PBS (control group). Competitive ELISA was performed in precoated 96-well plates, according to manufacturer’s instructions (Abbexa).

### Minimum inhibitory concentration assays

MIC assays were performed by broth microdilution. Isolates were first streaked onto fresh MH agar, and then a single colony from each plate was further grown overnight in 5 ml MH broth on an orbital shaker (180 r.p.m) at 37 °C. A fresh dilution in MH broth was made by incubating 200 µl of the overnight culture in 5 ml MH media. A hundred microliters of this culture was incubated in 96 well-plates with 100 µl of 1:2 serially diluted antibiotic in MH broth. After a 24 h static incubation at 37 °C, the OD_600_ was determined to assess bacterial growth.

### MIC assays with spermidine

MIC assays were performed as above, with the addition of 4 mM spermidine to the assay throughout or using *P. aeruginosa* that had been pre-incubated with 4 mM spermidine. In the latter case, overnight cultures of *P. aeruginosa*, prepared as above, were pelleted by centrifugation and resuspended in 5 ml PBS containing 4mM spermidine. Tubes were incubated at 37 °C, 180 r.p.m. for 30 minutes and then bacteria were again pelleted and resuspended in MH broth for use in MIC assays.

### Synthesis of fluorescent spermidine-NBD (nitrobenzoxadiazole)

7-Nitro-2,1,3-benzoxadiazol-4-N^8^-spermidine hydrogen chloride (NBD-N^8^-Spermidine)

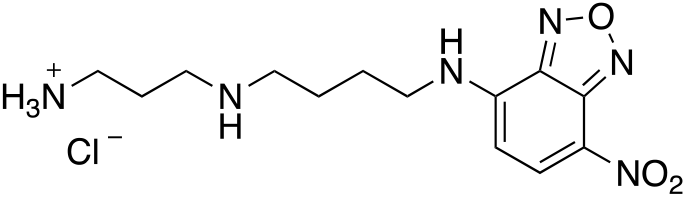

A solution of N^1^-N^4^-Bis-Boc-spermidine (50 mg, 0.29 mmol) in acetonitrile (2.0 ml) was added to 4-chloro-7-nitrobenzofurazan (58 mg, 0.29 mmol) and caesium carbonate (94 mg, 0.29 mmol) and heated to 80 °C under reflux conditions for 40 minutes. The reaction mixture was concentrated *in vacuo*, and purified through a silica plug, flushed with EtOAc to elute the boc protected fluorophore. The compound was then dissolved in a solution of 4M HCl in dioxane (5 mL) and stirred for 3 hours at room temperature, before removing the solvent *in vacuo*. The crude mixture was dissolved in DCM was and extracted with water. The aqueous layer was lyophilised to give NBD-N^8^-Spermidine as a hydrochloride salt (35) as a red, amorphous solid and a global yield of 74%, after both the addition of the fluorophore and the subsequent removal of the boc protection group. δ_H_ (500 MHz, D_2_O) 8.48 (1H, d, J 9.0, Ar 6-H), 6.35 (1H, d, J 9.0, Ar 5-H), 3.66 (2H, broad s, 2-H2), 3.17 (4H, m, 5- and 7-H2), 3.11 (2H, appt t, J 7.9, 9-H2), 2.08 (2H, apt tt, J 8, 2.4, 8-H2), 1.88 (4H, m, 3 and 4-H2). δ_C_ (125 MHz, D_2_O) 147.2 (Ar 4-C), 145.0 (Ar 7a-C), 144.7 (Ar 3a-C), 139.8 (Ar 5-C), 100.6 (Ar 6-C), 48.0 (5-C), 45.1 (7-C), 43.6 (2-C), 37.2 (9-C), 25.3 (3-C), 24.4 (8-C), 23.8 (4-C). HRMS (ESI): C_13_H_20_N_6_O_3_ requires [M+H]+, calculated 309.1675, found 309.1673.

### Flow cytometry analysis of spermidine-*P. aeruginosa* interactions

*P. aeruginosa* were pre-incubated with 4 mM spermidine-NBD or unlabelled spermidine, following the same protocol as that used for MIC assays. Following incubation, bacteria were pelleted by centrifugation and resuspended in PBS. Immediately, and at 30, 60, 120,180 and 240 minutes, 200 µl samples were removed from the cultures and analysed for NBD fluorescence on a FACS Aria II flow cytometer (BD Biosciences). Twenty thousand individual bacteria were recorded. Side-scatter and forward-scatter limits for bacterial flow cytometry were pre-determined using *P. aeruginosa* stained with the DNA dye thiazole orange (Sigma Aldrich).

### Biofilm assay

Biofilm experiments were set up using overnight cultures. Each culture was first diluted 1:100 in fresh broth, then added to 96 well plates and incubated for 48 hours at 37° C under static conditions. Plates were washed twice with PBS and stained with 200 μL of a 0.25% solution of crystal violet in water. After incubating at room temperature for 15 minutes, the plates were rinsed twice with water and allowed to dry for 24 hours. The stain was then dissolved in 1 mL of 25% acetic acid in water and incubated at room temperature for 2 minutes. Biofilm formation was quantified by measuring the optical density of this final solution at 590 nm.

### Statistics

Data analysis was carried out using GraphPad Prism v.8.02 and JMP version 14.0. Data were tested for normality. One-way or two-way ANOVA was used for comparison between groups, and post-hoc analysis included correction for multiple comparisons. Significance was determined from label free proteomics data using Progenesis QI.

### Study Approval

Ethical approval for collection of CF sputum was obtained from the North West Research Ethics Committee (IRAS:216408, ethics reference no: 17/NW/0091). Written informed consent was obtained from all study participants, prior to enrolment. Ethical approval for animal studies was obtained from the UK Home Office (project licence PP2072053) and the University of Liverpool Animal Welfare Ethical Review Board.

## Author Contributions

CW, AK, MW, DN and JF designed the study, contributed resources and reagents and supervised staff. CH, AG, AC, TJ, DN performed experiments. CH, AG, AC, MW, DN, JF analysed data. CH, DN, JF wrote the manuscript, with input from all authors.

## Acknowledgements

This work was funded by an Action Medical Research grant awarded to JF, DN, CW and AK (Grant number GN2444). AC was supported by an MRC Doctoral Training Partnership studentship. AG and DN were supported by a Wellcome and Royal Society Sir Henry Dale Fellowship awarded to DN (Grant Number 204457/Z/16/Z). JF was supported by a Medical Research Foundation Fellowship (Grant number MRF-091-0006-RG-FOTHE). JW was supported by a Leverhulme Trust Research Grant (Grant number RPG-2018-030).

## References

1. Rajan S, and Saiman L. Pulmonary infections in patients with cystic fibrosis. Semin Respir Infect. 2002;17(1):47–56.

2. Sadikot RT, Blackwell TS, Christman JW, and Prince AS. Pathogen-host interactions in Pseudomonas aeruginosa pneumonia. Am J Respir Crit Care Med. 2005;171(11):1209–23.

3. Lyczak JB, Cannon CL, and Pier GB. Lung infections associated with cystic fibrosis. Clin Microbiol Rev. 2002;15(2):194–222.

4. Langendonk RF, Neill DR, and Fothergill JL. The Building Blocks of Antimicrobial Resistance in Pseudomonas aeruginosa: Implications for Current Resistance-Breaking Therapies. Front Cell Infect Microbiol. 2021;11:665759.

5. Govan JR, and Deretic V. Microbial pathogenesis in cystic fibrosis: mucoid Pseudomonas aeruginosa and Burkholderia cepacia. Microbiol Rev. 1996;60(3):539–74.

6. Johnson L, Mulcahy H, Kanevets U, Shi Y, and Lewenza S. Surface-localized spermidine protects the Pseudomonas aeruginosa outer membrane from antibiotic treatment and oxidative stress. J Bacteriol. 2012;194(4):813–26.

7. Gunn JS. Bacterial modification of LPS and resistance to antimicrobial peptides. J Endotoxin Res. 2001;7(1):57–62.

8. Ernst RK, Moskowitz SM, Emerson JC, Kraig GM, Adams KN, Harvey MD, et al. Unique lipid a modifications in Pseudomonas aeruginosa isolated from the airways of patients with cystic fibrosis. J Infect Dis. 2007;196(7):1088–92.

9. Raetz CR, Reynolds CM, Trent MS, and Bishop RE. Lipid A modification systems in gram-negative bacteria. Annu Rev Biochem. 2007;76:295–329.

10. McPhee JB, Lewenza S, and Hancock RE. Cationic antimicrobial peptides activate a two-component regulatory system, PmrA-PmrB, that regulates resistance to polymyxin B and cationic antimicrobial peptides in Pseudomonas aeruginosa. Mol Microbiol. 2003;50(1):205–17.

11. McPhee JB, Bains M, Winsor G, Lewenza S, Kwasnicka A, Brazas MD, et al. Contribution of the PhoP-PhoQ and PmrA-PmrB two-component regulatory systems to Mg2+-induced gene regulation in Pseudomonas aeruginosa. J Bacteriol. 2006;188(11):3995–4006.

12. Mulcahy H, Charron-Mazenod L, and Lewenza S. Extracellular DNA chelates cations and induces antibiotic resistance in Pseudomonas aeruginosa biofilms. PLoS Pathog. 2008;4(11):e1000213.

13. Moskowitz SM, Brannon MK, Dasgupta N, Pier M, Sgambati N, Miller AK, et al. PmrB mutations promote polymyxin resistance of Pseudomonas aeruginosa isolated from colistin-treated cystic fibrosis patients. Antimicrob Agents Chemother. 2012;56(2):1019–30.

14. Moskowitz SM, Ernst RK, and Miller SI. PmrAB, a two-component regulatory system of Pseudomonas aeruginosa that modulates resistance to cationic antimicrobial peptides and addition of aminoarabinose to lipid A. J Bacteriol. 2004;186(2):575–9.

15. Han ML, Zhu Y, Creek DJ, Lin YW, Anderson D, Shen HH, et al. Alterations of Metabolic and Lipid Profiles in Polymyxin-Resistant Pseudomonas aeruginosa. Antimicrob Agents Chemother. 2018;62(6).

16. Winsor GL, Griffiths EJ, Lo R, Dhillon BK, Shay JA, and Brinkman FS. Enhanced annotations and features for comparing thousands of Pseudomonas genomes in the Pseudomonas genome database. Nucleic Acids Res. 2016;44(D1):D646–53.

17. Shah P, and Swiatlo E. A multifaceted role for polyamines in bacterial pathogens. Mol Microbiol. 2008;68(1):4–16.

18. Kwon DH, and Lu CD. Polyamine effects on antibiotic susceptibility in bacteria. Antimicrob Agents Chemother. 2007;51(6):2070–7.

19. Grasemann H, Shehnaz D, Enomoto M, Leadley M, Belik J, and Ratjen F. L-ornithine derived polyamines in cystic fibrosis airways. PLoS One. 2012;7(10):e46618.

20. Twomey KB, Alston M, An SQ, O’Connell OJ, McCarthy Y, Swarbreck D, et al. Microbiota and metabolite profiling reveal specific alterations in bacterial community structure and environment in the cystic fibrosis airway during exacerbation. PLoS One. 2013;8(12):e82432.

21. Fothergill JL, Neill DR, Loman N, Winstanley C, and Kadioglu A. Pseudomonas aeruginosa adaptation in the nasopharyngeal reservoir leads to migration and persistence in the lungs. Nat Commun. 2014;5:4780.

22. Bricio-Moreno L, Sheridan V, Goodhead IB, Armstrong S, Wong JKL, Waters EM, et al. Evolutionary trade-offs associated with loss of PmrB function in host-adapted Pseudomonas aeruginosa. Nat Commun. 2018.

23. Bolard A, Schniederjans M, Haussler S, Triponney P, Valot B, Plesiat P, et al. Production of Norspermidine Contributes to Aminoglycoside Resistance in pmrAB Mutants of Pseudomonas aeruginosa. Antimicrob Agents Chemother. 2019;63(10).

24. Karatan E, and Michael AJ. A wider role for polyamines in biofilm formation. Biotechnol Lett. 2013;35(11):1715–7.

25. McGinnis MW, Parker ZM, Walter NE, Rutkovsky AC, Cartaya-Marin C, and Karatan E. Spermidine regulates Vibrio cholerae biofilm formation via transport and signaling pathways. FEMS Microbiol Lett. 2009;299(2):166–74.

26. Goytia M, Dhulipala VL, and Shafer WM. Spermine impairs biofilm formation by Neisseria gonorrhoeae. FEMS Microbiol Lett. 2013;343(1):64–9.

27. Moore MP, Lamont IL, Williams D, Paterson S, Kukavica-Ibrulj I, Tucker NP, et al. Transmission, adaptation and geographical spread of the Pseudomonas aeruginosa Liverpool epidemic strain. Microb Genom. 2021;7(3).

28. Bricio-Moreno L, Sheridan VH, Goodhead I, Armstrong S, Wong JKL, Waters EM, et al. Evolutionary trade-offs associated with loss of PmrB function in host-adapted Pseudomonas aeruginosa. Nat Commun. 2018;9(1):2635.

29. Durand JM, and Bjork GR. Putrescine or a combination of methionine and arginine restores virulence gene expression in a tRNA modification-deficient mutant of Shigella flexneri: a possible role in adaptation of virulence. Mol Microbiol. 2003;47(2):519–27.

30. Zhou L, Wang J, and Zhang LH. Modulation of bacterial Type III secretion system by a spermidine transporter dependent signaling pathway. PLoS One. 2007;2(12):e1291.

31. Delcour AH. Outer membrane permeability and antibiotic resistance. Biochim Biophys Acta. 2009;1794(5):808–16.

32. Bonfield TL. Preclinical Modeling for Therapeutic Development in Cystic Fibrosis. Am J Respir Crit Care Med. 2020;201(3):267–8.

33. Moyne O, Castelli F, Bicout DJ, Boccard J, Camara B, Cournoyer B, et al. Metabotypes of Pseudomonas aeruginosa Correlate with Antibiotic Resistance, Virulence and Clinical Outcome in Cystic Fibrosis Chronic Infections. Metabolites. 2021;11(2).

34. Cornforth DM, Diggle FL, Melvin JA, Bomberger JM, and Whiteley M. Quantitative Framework for Model Evaluation in Microbiology Research Using Pseudomonas aeruginosa and Cystic Fibrosis Infection as a Test Case. mBio. 2020;11(1).

35. Jagu E, Pomel S, Pethe S, Cintrat JC, Loiseau PM, and Labruere R. Spermine-NBD as fluorescent probe for studies of the polyamine transport system in Leishmania donovani. Bioorg Med Chem Lett. 2019;29(14):1710–3.

