## Supplementary Figures for "*Pseudomonas aeruginosa* utilises host-derived polyamines to facilitate antimicrobial tolerance"

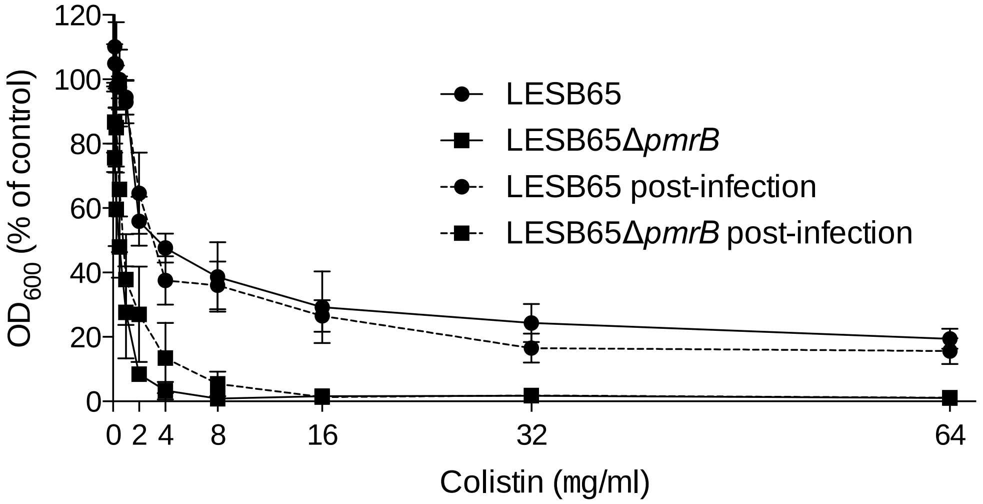


**Supplementary Figure 1. Colistin sensitivity of LESB65 and LESB65Δ*pmrB* before and after two-day infection of mouse airways.** Post-infection samples are *P. aeruginosa* recovered from 2 day post-infection mouse lung homogenates. Bacteria were recovered by plating homogenates onto Pseudomonas selective agar and then growth for ~30 hours at 37°C. Susceptibility testing was performed by microdilution in Mueller Hinton broth with two-fold decreasing concentrations of colistin (range 64-0.125µg/ml). Post-infection samples are from a single *in vivo* experiment. Frozen stocks of these bacteria were tested for susceptibility on three separate assay days. Data show mean $OD_{600}$ ± standard deviation.

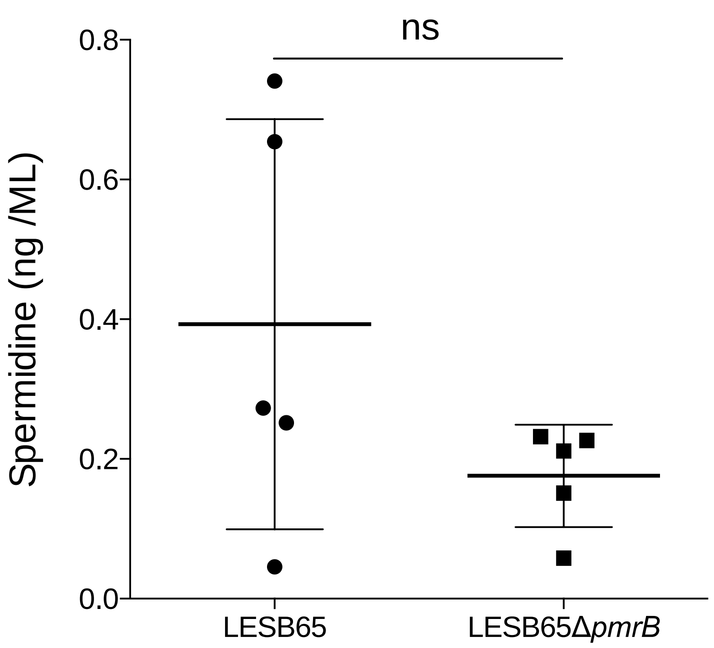


**Supplementary Figure 2. Spermidine levels in bacterial lysates.** Lysates of overnight cultures were used in a competitive spermidine ELISA. Levels detected were close to the manufacturer's stated limits of assay sensitivity (<0.28 ng/ml). Data are from five separate overnight cultures per strain, run, in duplicate, on a single ELISA assay plate. Ns = not significant.

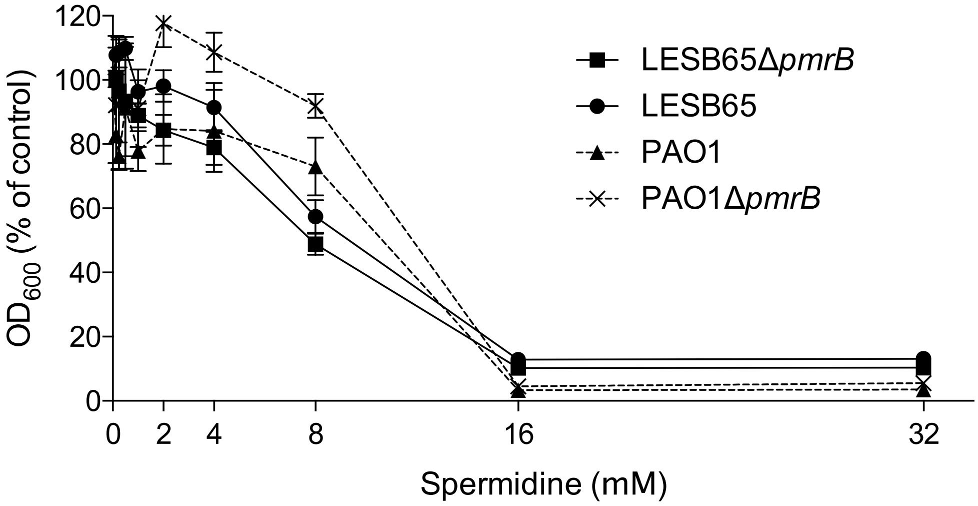


**Supplementary Figure 3. Growth of *P. aeruginosa* in the presence of exogenous spermidine.** LESB65, PAO1 and their isogenic *pmrB*-deficient mutants were grown in Mueller Hinton broth in the presence of spermidine (range 0.125-32 mM). Growth was determined after 24 hours by reading absorbance at 600 nM. A concentration of 4 mM was chosen for subsequent assays.

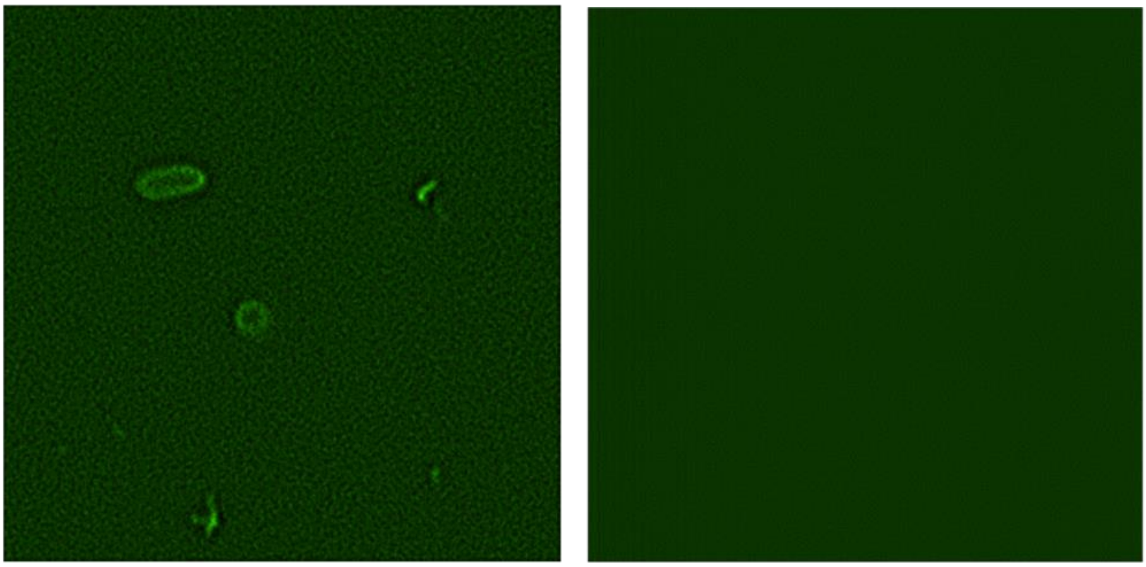

**Supplementary Figure 4. Spermidine-NBD coats the *P. aeruginosa* surface.** Fluorescence microscopy images of LESB65 in the presence (left) or absence (right) of spermidine-NBD.
